# How size and shape affect the vertical velocity of cyanobacterial colonies

**DOI:** 10.64898/2026.03.25.714148

**Authors:** Yuri Z. Sinzato, Jolanda M. H. Verspagen, Robert Uittenbogaard, Petra M. Visser, Jef Huisman, Maziyar Jalaal

## Abstract

Cyanobacterial colonies often exploit their buoyancy and large size to float upwards rapidly and form dense surface blooms, which can degrade water quality, threaten ecosystems and public health, and impose substantial economic costs. Yet, how the morphology of cyanobacterial colonies controls their vertical velocity remains poorly understood. We conducted detailed three-dimensional morphological characterization of colonies of the cyanobacterium *Microcystis* in lake samples at the single-colony level and performed controlled flotation experiments in stratified flows. Using particle tracking in a vertical density gradient, we separately quantified the contributions of colony shape and buoyant density at the level of individual colonies. Our results show that the shape factor in Stokes’ law varies systematically with colony size. Consequently, the vertical velocity of colonies does not scale with the square of colony size but only with a power of 1.13, as larger colonies have a more irregular shape and therefore experience enhanced drag. We therefore correct the commonly used Stokes’ law to account for the size-dependent change in the shape factor. Interestingly, implementation of this power law relationship in a vertical migration model shows widespread chaotic dynamics in the migration trajectories of *Microcystis* colonies. These results highlight the importance of morphological plasticity in cyanobacterial colonies and can inform predictive models and hydrodynamic control strategies for toxic blooms. Our methodology to simultaneously determine the density, shape factor and velocity is broadly applicable to suspended aggregates with complex shapes in freshwater and marine systems.

## 1. INTRODUCTION

Cyanobacterial blooms in lakes and reservoirs can cause fish kills, foul water supplies, and expose humans, livestock, and wildlife to hazardous toxins [1, 2]. One of the most common bloom-forming taxa is *Microcystis*, a globally prevalent genus of both freshwater and slightly brackish ecosystems whose large, buoyant colonies accumulate near the surface and drive recurrent toxic events [3]. The vertical motion of *Microcystis* is a key trait behind the widespread success of this cyanobacterium [4]. Large buoyant colonies in non-turbulent water have better access to light compared to non-buoyant algae due to high flotation velocities [5], while also being able to escape regions of high light intensity via a density regulation mechanism [6, 7]. Hence, understanding the factors that influence the vertical velocity of *Microcystis* colonies is essential for hydrodynamic-based prediction and control techniques. In fact, models have been developed to predict the vertical migration of colonies, based on the balance between synthesis and consumption of carbohydrates [7–11]. These migration models can ultimately be coupled to hydrodynamic equations to predict the growth and spatial distribution of *Microcystis* populations in wind-mixed lakes [12–14]. Furthermore, in-lake controls such as artificial deep mixing have been used, with partial success, to increase turbulence and counteract the buoyant advantage of *Microcystis* colonies [15]. Here too, precise estimates of the flotation velocity are essential for the design of the mixing system, as weak turbulence may not overcome the flotation of colonies and thus fail to suppress cyanobacterial growth.

In a regime of negligible fluid inertia, colonies in a uniform and quiescent water column move with a vertical velocity *U* described by Stokes’ law [16],

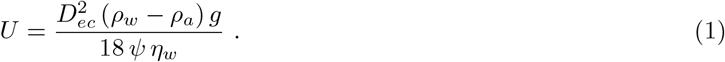

Here *D*_*ec*_, *ρ*_*a*_ and *ψ* are the equivalent circular diameter, the density and the shape factor of the colony, respectively, while *ρ*_*w*_ and *η*_*w*_ are the density and dynamic viscosity of water, respectively, and *g* is the acceleration of gravity. Eq. (1) results from the balance between the net buoyancy force and viscous drag of the colony (*Supporting Information Text* S1). *Microcystis* colonies are made up of spherical cells joined together by a transparent mucilage layer, composed of extracellular polymeric substances [17, 18]. Despite lacking active motility, these colonies move vertically by adjusting their density. The balance between intracellular gas vesicles (lowering density) and carbohydrate ballast (raising density) sets a time-dependent colony density *ρ*_*a*_(*t*) that alternates between positive and negative buoyancy [7, 19]. The mucilage layer contributes to the total colony volume, while its density is similar to that of water, thus the mucilage attenuates the amplitude of colony density fluctuations [20].

Beyond buoyancy, colony shape also controls its vertical velocity. In Eq. (1), the influence of colony shape on velocity is quantified with a shape factor *ψ*; where a larger *ψ* denotes a greater departure from sphericity, which increases drag and thereby reduces vertical velocity. The classic Stokes’ law can be recovered from Eq. (1) by setting the shape factor to *ψ* = 1, and is valid for spherical particles. A common assumption in several hydrodynamic models for the cyanobacterial colonies is that the vertical velocity follows the classic Stokes’ law, where the colony velocity is proportional to the square of its size [13, 14, 21]. However, Nakamura *et al*. [22] have previously demonstrated that the approximation of spherical shape is invalid for *Microcystis* colonies, because their shape varies with their size. The shape–size dependence can be captured by a fractal relation, 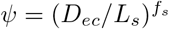, valid for *D*_*ec*_≥ *L*_*s*_, where *f*_*s*_ is the shape fractal exponent and *L*_*s*_ is a characteristic size above which colonies deviate from the spherical geometry. Applying the fractal relation for the shape factor to Eq. (1) results in a velocity that scales with the colony size to a power (i.e., 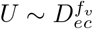), where *f*_*v*_ = 2 − *f*_*s*_. Nakamura *et al*. [22] found a power of *f*_*v*_ = 1.5 for *Microcystis* colonies sampled from Lake Kasumigaura (Japan). Li *et al. [23]* proposed an alternative shape–size dependence given by a second-order polynomial, where the measured coefficients varied for different morphospecies of *Microcystis* colonies sampled from Lake Taihu (China).

The diversity of *Microcystis* morphospecies and the broad range of colony sizes complicate high-fidelity 3D imaging and quantitative shape characterization of *Microcystis* colonies, and by extension, their hydrodynamic characteristics [4, 24]. Prior studies have measured single-colony flotation velocities in uniform water columns using particle tracking [22, 23, 25, 26]. However, direct measurements of the buoyant density (i.e., the colony density in its buoyant state) of single colonies are lacking. Instead, buoyant density has typically been inferred by applying Stokes’ law to measured flotation velocities [23, 26, 27], an approach that relies on strong assumptions about the colony shape factor. Alternatively, the density of floating colonies has been estimated by separately measuring intracellular gas volume and the density of colonies after gas-vesicle collapse [28]. This method yields only suspension-averaged values and cannot resolve single-colony densities. Measurements in a uniform-density column are also vulnerable to thermal convection, which introduces significant measurement noise for slowly rising or sinking particles [29]. To suppress convection, a continuous density gradient can be imposed [30, 31]. However, such gradients alter the hydrodynamics: rising (sinking) particles entrain denser (lighter) fluid, adding resistance and biasing velocity and density estimates [32–34]. These limitations have impeded the development of a single-colony–resolved, mechanistic parameterization for vertical migration and, consequently, have constrained bloom forecasting and the design of effective in-lake control strategies. Hence, here, we pose a central question: can we directly measure, at single-colony resolution, both the density and shape factor of *Microcystis* colonies to obtain independent estimates of their respective contributions to the size-dependent vertical velocity of these colonies in natural populations?

To address this question, we quantified the vertical motion of individual *Microcystis* spp. colonies sampled from Lake Braassemermeer and Lake’t Joppe in the Netherlands. Both lakes are used for recreation and suffer from recurrent cyanobacterial blooms (Fig. 1). We integrated optical coherence tomography (OCT), confocal fluorescence microscopy, mucilage staining, and gas-fraction measurements to resolve meso- and microscale colony structures and corrected for the projection-bias in colony volume estimates. Single colonies were measured and tracked in a thermally stable linear density gradient; from each trajectory we simultaneously inferred the shape factor *ψ* and the colony density *ρ*_*a*_. Hence, we obtained a large dataset with which we assessed the relationship between the flotation velocity and the size, shape and density of individual *Microcystis* colonies. Lastly, we applied the experimental results to a vertical migration model to showcase how the size-dependent shape factor affects the migration patterns of *Microcystis* colonies. Beyond advancing basic understanding of bloom-forming cyanobacteria, our analysis provides parameters and measurements needed for predictive transport models of algal colonies and for designing in-lake control strategies of cyanobacterial blooms.

**Fig. 1:**
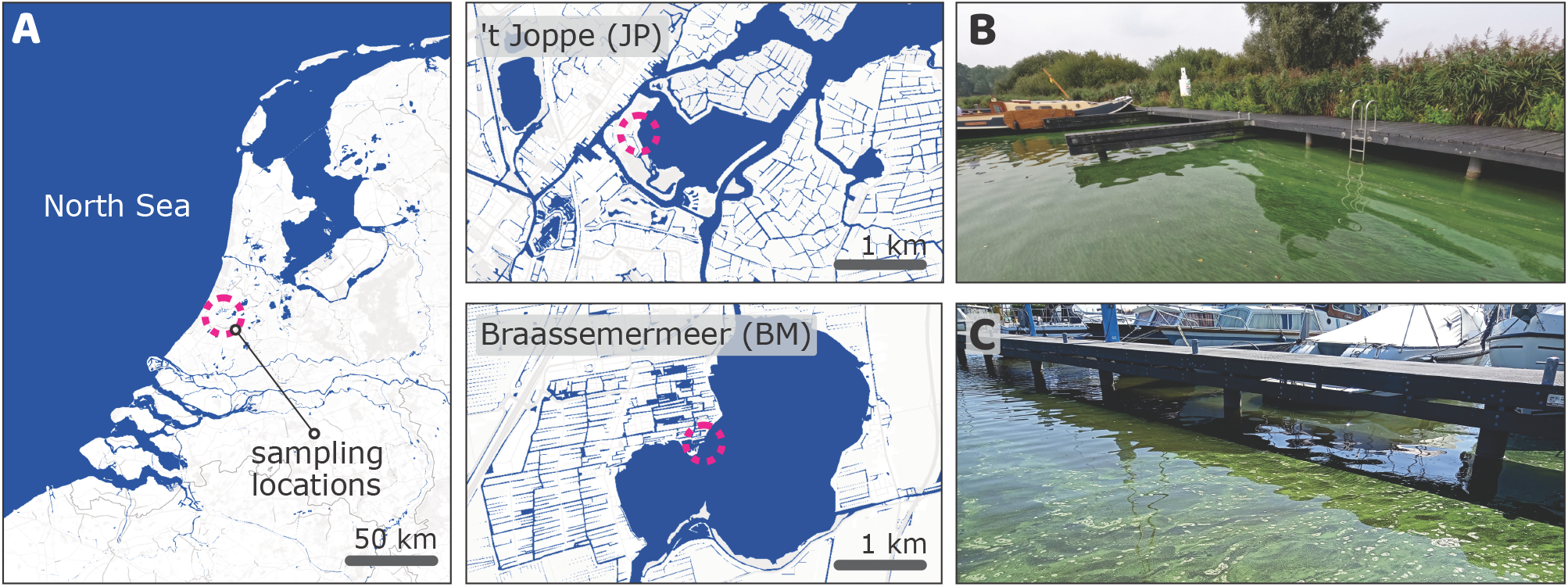
(A) Map of the sampled lakes with the sampling locations circled. *Microcystis* spp. colonies were collected from the surface layer of (B) Lake’t Joppe and (C) Lake Braassemermeer.

## 2. MATERIALS AND METHODS

### 2.1. Samples

*Microcystis* spp. colonies were collected from Lake Braassemermeer (6^th^ August 2024) and Lake’t Joppe (3^rd^ September 2024), in the Netherlands. Plankton samples were concentrated with a 35 µm net and transported to the laboratory in cooled vials within 3 hours of collection. Upon arrival, the samples were centrifuged at 100 g for 15 minutes to separate buoyant cyanobacteria from other non-buoyant plankton. The abundance of *Microcystis* spp. colonies [35] in the buoyant fraction was above 98 % for both lakes (*N* = 101 counted colonies for Lake’t Joppe and *N* = 143 for Lake Braassemermeer), with *M. ichthyoblabe* being the most abundant morphospecies (45 % for Lake’t Joppe and 69 % in Lake Braassemermeer), followed by *M. aeruginosa* (44 % and 17 % respectively) and *M. flos-aquae* (14 % and 10 % respectively). A subsample of the buoyant fraction was separated for measurement of its gas vesicle content, while the remaining sample was incubated for 24 hours, in the dark, at 20 °C, under gentle aeration. This dark incubation promoted the consumption of accumulated carbohydrates, and the duration was sufficiently long to reduce the density of an initially buoyant colony to its minimum value [10]. After incubation, measurements of flotation velocity were conducted within an hour, while morphological analysis was performed within 3 days on refrigerated samples (dark, 4 °C).

### 2.2. Morphological analysis

The morphology of *Microcystis* colonies is composed of mesoscale features at the colonial scale (∼ 10^2^ µm to 10^3^ µm) and microscale features at the cellular scale (∼ 1 µm to 10 µm). In this section, we first describe the main morphological descriptors associated with *Microcystis* colonies and later describe the imaging methods used to measure the morphological features. The buoyancy force experienced by a *Microcystis* colony depends on its volume (*Supporting Information Text* S1). However, two-dimensional projections from transmitted-light microscopy cannot resolve the structure along the projection (optical) axis. We therefore used an OCT imaging system to characterize the 3-dimensional mesoscale features of *Microcystis* colonies. At the microscale, below the OCT resolution, mucilage staining, confocal microscopy, gas vesicle fraction measurements, and cell diameter measurements were performed to measure morphological features.

#### 2.2.1. Morphological descriptors for *Microcystis* colonies

Due to the non-homogeneous and irregular structure of *Microcystis* colonies, a single size descriptor cannot uniquely represent it. Instead, the suitable characteristic size *L*_*c*_ will depend on the intended use and the available measuring techniques. We classify the colony structure into cellular and mucilage components, such that the total colony volume is *V*_*c*_ = *V*_*m*_ + *N*_*c*_ *V*_1_, where *V*_*m*_ is the volume of the mucilage layer, *N*_*c*_ is the number of cells inside a colony, and *V*_1_ is the single cell volume, which is assumed to be constant throughout the colony. Liquid regions that fill voids or holes between mucilage regions are not considered part of the colony. The colony volume can be used to define a characteristic size, named equivalent spherical diameter *D*_*es*_, such that *D*_*es*_ = (6 *V*_*c*_*/π*)^1/3^. However, in a two-dimensional imaging system, only the projected area *A*_*c*_ of the colony along the imaging axis is available. In this case, one may use the equivalent circular diameter *D*_*ec*_ = (4 *A*_*c*_*/π*)^1/2^ as the characteristic size. We define the ratio between the colony volume and the projection-based volume as the volume factor *ζ* = (*D*_*es*_*/D*_*ec*_)^3^. Although *D*_*ec*_ = *D*_*es*_ (*i*.*e*., *ζ* = 1) for solid spheres, these two characteristic sizes will deviate for more complex morphologies [36]. A fractal representation, *i*.*e*., a power law dependence, can be used to relate the volume factor and the equivalent circular diameters of *Microcystis* colonies, 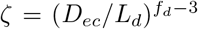, valid for *D*_*ec*_ ≥*L*_*d*_ [37]. Consequently, the equivalent spherical and circular diameters are related by 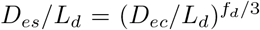. Here, *L*_*d*_ is the characteristic size above which the colony departs from a homogeneous spherical shape, and *f*_*d*_ is the fractal dimension of the colony.

The colony density *ρ*_*a*_ is a weighted average of the cell and mucilage densities, *ρ*_*c*_ and *ρ*_*m*_ respectively, such that *ρ*_*a*_ = *ρ*_*c*_ *ϕ* + *ρ*_*m*_ (1 − *ϕ*), where *ϕ* is cell-to-colony volume fraction. Each cell contains on average a volume *V*_*g*_ of gas in gas vesicles, such that the per–cell gas fraction is *ϕ*_*g,c*_ = *V*_*g*_*/ V*_1_. When subjected to high pressure, the gas vesicles collapse, and water enters the cell to partially replace the gas volume *V*_*g*_ [28]. The result is a change in the cell density that can be approximated by Δ *ρ*_*c*_ = *ϕ*_*g,c*_ *ρ*_*w*_ (*Supporting Information Text* S3). When considering the total colony volume (cell+mucilage), the change in colony density after gas vesicle collapse is Δ *ρ*_*a*_ = *ϕ*_*g,a*_ *ρ*_*w*_, where *ϕ*_*g,a*_ is the per-colony gas fraction. Since the density of mucilage is similar to that of water [20], the two definitions of gas fraction are related by *ϕ* = *ϕ*_*g,a*_*/ ϕ*_*g,c*_. We summarize the morphological descriptors for cyanobacterial colonies in the Supplementary Table S1.

#### 2.2.2. Optical coherence tomography

An OCT imaging system (Telesto TEL221C1(/M) unit equipped with a LSM04 lens, Thorlabs) was used to characterize the mesoscale structure of field-collected *Microcystis* spp. colonies from Lake’t Joppe. OCT is a non-invasive, low-coherence interferometric imaging technique that reconstructs volumetric data with micrometer-scale axial resolution in weakly scattering samples [38]. Gas vesicles were collapsed with a pressure of 1 MPa of compressed nitrogen gas and the colonies were injected between microscope slides with a 350 µm spacer. The microscope slide was positioned at an angle of 5° relative to the OCT scanning axis to reduce reflection from the glass surfaces. The field of view allows for a 3-dimensional scan of the entire colony. The three-dimensional scans were subjected to noise reduction with Gaussian blur, thresholding, and connection of nearly touching object regions, after which the colonies were segmented and measured using the Python library `scikit-image` [39]. The lateral resolution of 20 µm was larger than individual cells. We hypothesized that the colony volume measured with OCT corresponds to the total colony volume *V*_*c*_, consisting of combined cells and mucilage, and we verified this hypothesis using mucilage staining and 3-dimensional confocal microscopy. Moreover, only colonies larger than 180 µm were imaged, given the lateral resolution of the OCT.

#### 2.2.3. Gas fraction and cell diameter measurements

The absolute gas-volume per cell *V*_*g*_ and the associated per–cell gas fraction *ϕ*_*g,c*_ of *Microcystis* spp. colonies was measured using a capillary compression tube [28]. The colonies were collected in a 10 µm nylon mesh and redispersed in sucrose solution (7 g/mL), in order to fragment large colonial aggregates. The sample was degassed in a dark vacuum jar for 5 minutes at 8 kPa. The suspension was injected into the capillary compression tube at a controlled temperature of 20 °C. The gas content in the suspension was measured from its volume reduction after application and release of 1 MPa pressure. Gas content measurements were taken in triplicates. The collapsed suspensions were collected for measurement of the cell concentration and the average cell diameter. Individual cells were imaged in a wide-field fluorescence microscope using chlorophyll *a* excitation and their diameter *d*_1_ was measured with an image processing script. Cells were assumed to be spherical, such that 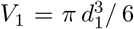. To measure the cell concentration in the suspension, the colonies were disaggregated into single cells via a heating treatment [40]. The suspension was injected into a Neubauer haemocytometer (Marienfeld) and the number of cells was counted with an image processing script.

#### 2.2.4. Mucilage visualization

The mucilage layer encasing the cells of *Microcystis* spp. colonies was stained with Alcian Blue 8GX (0.3 % (w/v) in acetic acid at 3 % (v/v)) [41]. Equal parts of the stain solution and colony suspension were mixed and incubated for 1 hour. To remove excess stain, a pressure of 1 MPa was applied to collapse the gas vesicles, colonies were allowed to sink, the supernatant was removed and colonies were re-dispersed in acetic acid at 3 % (v/v). The stained colonies were injected between microscope slides with a 350 µm spacer. The colonies were imaged with brightfield microscopy, and the blue and red channels were captured with a DAPI (emission between 435 and 485 nm, Nikon) and a Cy5 (emission between 672 and 712 nm, Edmund Optics) band-pass filters, respectively. The images were normalized for background intensity and the difference between the blue and red channels was calculated. The stained mucilage layer appeared as bright regions, since it transmitted the blue light while absorbing the red light.

#### 2.2.5. Confocal microscopy

The 3-dimensional arrangement of cells within a colony was investigated with laser scanning confocal microscopy. Gas vesicles were collapsed with a pressure of 1 MPa of compressed nitrogen gas and the colonies were injected between microscope slides with a 350 µm spacer. The chlorophyll *a* auto-fluorescence (excitation at 636 nm) was used for visualization of the cells. Only the first few cell layers were observable, due to the light attenuation at deeper layers. Cells were segmented using StarDist [42]. Cell centroids were measured, from which the nearest-neighbor distance (NND) for each cell was calculated with an image processing script. The nearest neighbor distances were normalized by the average cell diameter *d*_1_ and the resulting distribution, 𝒫 (NND*/ d*_1_), was computed.

### 2.3. Flotation velocity measurements

#### 2.3.1. Flotation chamber

Flotation velocity measurements were conducted in a density gradient of Percoll (Cytiva) (Fig. 2A). Percoll was used as it forms stable gradients with low viscosity, low diffusivity, and minimal osmotic effects. The inner dimensions of the chamber are 80 mm × 27 mm × 27 mm. Two opposing glass windows allowed for back illumination from a white LED panel and optical access to the cameras. The temperature in the liquid column was regulated at 20 °C (±0.05 °C) via two lateral chambers with water circulation. The column was filled using the standard two-cylinder method [43]. A cylinder filled with standard BG-11 medium [44] flowed by gravity into a second cylinder with the same volume of a denser solution (22 % (w/w) of Percoll and 2 % (w/w) of 50x BG-11 medium). The liquids were mixed in the latter cylinder and slowly flowed into the measuring chamber via a long needle. The density *ρ*_*l*_ and dynamic viscosity *η*_*l*_ of the liquid in the chamber varied linearly with the height *z* (measured from the chamber bottom), such that *ρ*_*l*_(*z*) = *ρ*_*o*_ − *β z* and *η*_*l*_(*z*) = *η*_*o*_ − *αz*. Here, *ρ*_*o*_ = 1.025 g/mL (± 0.0006 g/mL) and *η*_*o*_ = 1.138 mPa s (± 0.003 mPa s) were the density and viscosity at the bottom, and *β* =0.35 m^*−*1^ g/mL (±0.02 m^*−*1^ g/mL) and *α* =1.76 m^*−*1^ mPa s (± 0.08 m^*−*1^ mPa s) were the density and viscosity gradients, respectively. The density and viscosity of the bulk solutions were measured with the gravimetric method and the rotational rheometry method, respectively [45].

**Fig. 2:**
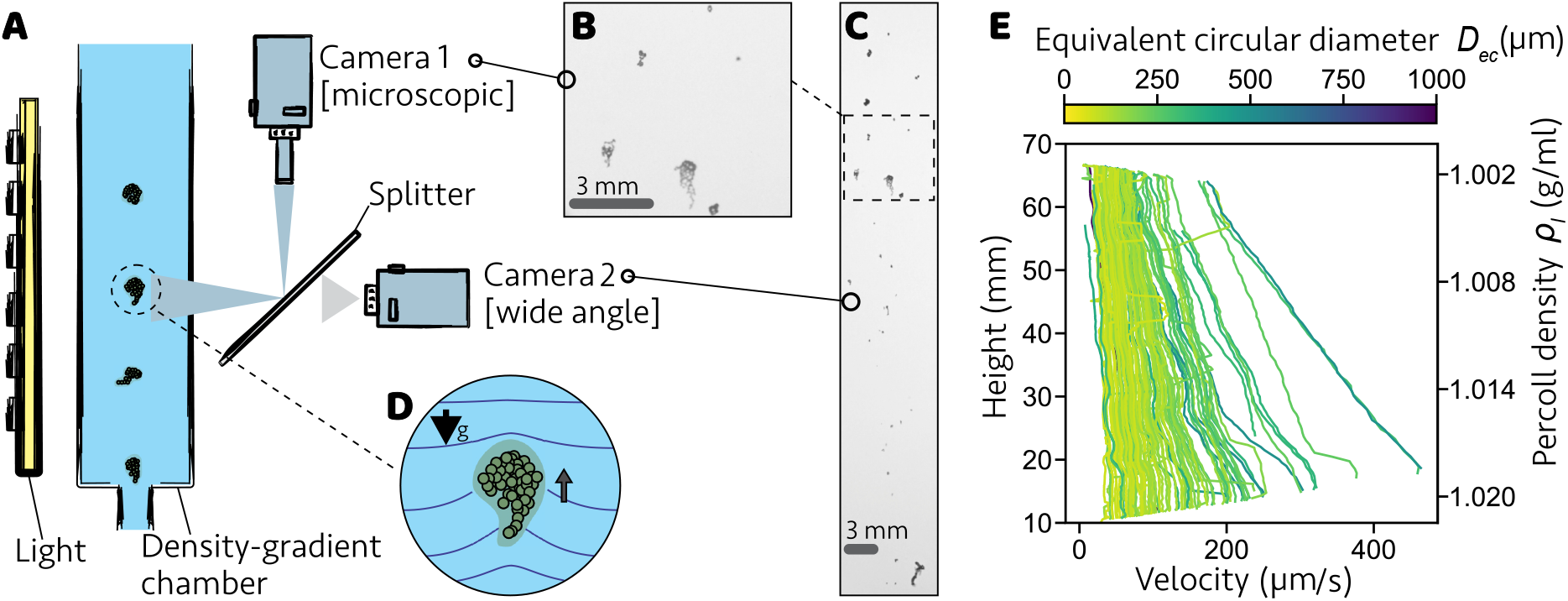
Experimental setup used for measuring the flotation velocity. (A) Buoyant colonies entered a liquid chamber filled with a density gradient of Percoll. The size and shape of each individual colony were measured from (B) the microscopic image, while its trajectory through the density gradient was visualized with (C) the wide-angle image. (D) As a colony rose, it produced a disturbance of the isopycnic lines (blue lines). (E) The rising velocity of each colony was measured as a function of its height from the chamber bottom and the associated Percoll density (*N* = 75 colonies from Lake Braassemermeer).

#### 2.3.2. Image acquisition

After the dark incubation described in section 2.1, the suspension was separated in two aliquots: one subsample for measurement of the shape factor and colony density with intact gas vesicles, and another subsample for the measurement of the colony density after collapse of gas vesicles. The first subsample was mixed with a dense solution (50 % (w/w) of Percoll and 2 % (w/w) of 50x BG-11 medium). The suspension of colonies was gently withdrawn with a syringe, which was subsequently connected to the bottom valve of the measuring chamber. The valve was opened and the colonies floated into the liquid column without fluid injection, in order not to disturb the flow field. The trajectory of the colonies through 70 % of the column height was imaged at a wide-angle view with a DSLR camera (Z6II, Nikon) (Fig. 2C). In addition, the light path was split in a microscopic image captured with a stereomicroscope (SMZ18, Nikon) connected to a second DSLR camera (D5200, Nikon) (Fig. 2B). The microscopic image was used for colony size and shape measurement. A calibrated target was used for pixel size determination and matching of both images. Only colonies larger than 50 µm were tracked, due to the lateral resolution (5 µm) of the microscopic image. The second subsample of colonies was subjected to 1 MPa pressure to collapse the gas vesicles. The resulting sinking colonies were gently withdrawn with a syringe and allowed to settle into the chamber via the upper valve. After 30 minutes, all colonies had reached their isopycnic layer. The density of each colony was extracted from their equilibrium height. The collapse of gas vesicles does not affect the structure of the colony, therefore it allowed us to vary the colony density without modifying its morphology.

#### 2.3.3. Colony tracking

The images underwent pre-processing with background subtraction, noise reduction with Gaussian blur, thresholding, and connection of nearly touching object regions using the Python library `scikit-image` [39]. For each segmented colony, its Feret diameter (maximum linear dimension), equivalent circular diameter, aspect ratio (major/minor axis), and orientation (mean angle between each colony’s major axis and its velocity direction) were measured using `scikit-image`. Colony trajectory linking was performed with the Python library trackpy [46]. A moving average was applied to the colony velocity trajectory to reduce noise. As the colony rises in the density gradient, its flotation velocity *U* (*z*) will decrease as the density excess, *ρ*_*l*_(*z*) − *ρ*_*a*_, decreases (Fig. 2E). An implicit expression for the flotation velocity *U* (*z*) can be obtained from the force balance on the colony (*Supporting Information Text* S1):

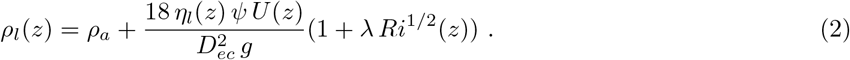

Here, *Ri*(*z*) = *β g* (*ψ ζ D*_*ec*_)^3^/ 8 *η*_*l*_(*z*) *U* (*z*) is the viscous Richardson number, which corrects the drag force on the colony for the entrainment of high density fluid [33] (Fig. 2D). For each floating colony, its velocity as a function of height was fitted with Eq. (2) using the shape factor *ψ* and the colony density *ρ*_*a*_ as fitting parameters (Fig. S1). The average orientation of the colonies was aligned with their velocity (mean angle ±SD between each colony’s major axis and its velocity direction was 1°± 33°, with *N* = 167 colonies; Fig. S2), indicating that, as in the OCT scans, the image projection axis was perpendicular to the colony’s major axis.

#### 2.3.4. Validation with microspheres

The methodology for the flotation velocity measurements was verified with calibrated microspheres (FluoRed PE, Cospheric) with a similar size and density as the colonies. Microspheres with a density between 0.9983 g/ml to 1.0003 g/ml (± 0.0006 g/ml) were fractioned in a two-layer column of different glycerol and water mixtures. A diameter of 148 ±5 µm (SD, *N* = 56) was measured with brightfield microscopy. The flotation velocity of the particles was well predicted by Eq. (2) within the instrumental uncertainty range (Fig. S3).

#### 2.3.5. Comparison of the flotation velocity measured in a discrete double layer and in a linear density gradient

Further verification of the methodology for the flotation velocity measurements was conducted with an alternative liquid column with a discrete double layer of Percoll (Fig. S4, (*Supporting Information Text* S2)). Two Percoll suspensions of different densities were layered inside the flotation chamber, producing two homogeneous density layers with a sharp interface (with respect to the chamber height). Velocity trajectories in a discrete double layer were measured for a subsample of buoyant colonies from Lake’t Joppe, following the same protocol for image acquisition described in section 2.3.2. The velocity greatly decreased near the sharp interface due to the fluid entrainment [47], therefore this region was excluded for the calculations of the hydrodynamic parameters. The velocity of each colony in the bulk of each layer was substituted in Eq. (2), with *Ri*(*z*) = 0 and the flotation velocity of the colony in water was computed (Fig. S4). The flotation velocities measured in a discrete double layer agree well with those measured in a linear density gradient (One-way ANCOVA, *Supporting Information Text* S2), with the former methodology being less suitable due to the large number of trajectories lost out of the field of view.

### 2.4. Vertical migration model

To assess how the size dependence of the shape factor will impact vertical migration of *Microcystis*, we applied a standard vertical migration model [6, 7, 10] to simulate migration trajectories of cyanobacterial colonies using Stokes’ law (1) with either (i) a constant shape factor *ψ* = 1 for all colony sizes or (ii) a size-dependent shape factor 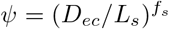 for *D*_*ec*_ ≥*L*_*s*_, and *ψ* = 1 for *D*_*ec*_ *< L*_*s*_. In this model, changes in colony density depend on the rate at which carbohydrates are produced by photosynthesis in the light and consumed by respiration in the dark, which results in vertical migration of the colonies. The model consists of a pair of ordinary differential equations describing changes in colony depth, and carbohydrate content (and consequently colony density) as a function of time. The model was parameterized based on the recent study of Feng *et al*. [10] and the values for all parameters are listed in the Supplementary Table S2. Values for shape factor parameters *L*_*s*_ and *f*_*s*_, cell-to-colony volume fraction *ϕ*, and per-cell gas fraction *ϕ*_*g,c*_ were estimated from the experiments in this study. Moreover, the modeled colony density at the minimum carbohydrate content was set to the average colony density estimated in this study. All other parameters are based on Feng *et al*. [10]. Vertical migration patterns were simulated for colonies with diameters ranging from 10 µm to 1000 µm over a period of 250 days but only analyzed from day 50 onward to remove initial transients and allow the migration dynamics to synchronize with the day-night cycle.

## 3. RESULTS AND DISCUSSION

### 3.1. Micro- and meso-scale characteristics of the colonies and the mismatch between the volume and projection-based sizes

OCT images of the three-dimensional mesoscale structure of field-collected *Microcystis* spp. colonies showed that large colonies often exhibited branched architectures and internal tunnels that were not visible in two-dimensional projections, as illustrated here for a typical *Microcystis aeruginosa* colony (Fig. 3A). Notably, the equivalent circular diameter of the colony, *D*_*ec*_, systematically overestimated the equivalent spherical diameter, *D*_*es*_, reflecting the loss of sub-surface structure in 2D projections (Fig. 3B). The two descriptors were linked by a power law, 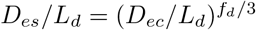, with *L*_*d*_ = 77 ± 20 µm and *f*_*d*_ = 2.36 ± 0.10 (*N* = 66 colonies, *R*^2^ = 0.89, dashed line in Fig. 3B). This geometric bias in *D*_*ec*_ propagates directly to buoyant force estimates, which scale with the total colony volume *V*_*c*_ (*Supporting Information Text* S1). Accordingly, when inferring buoyant force from projected area alone, a volume correction factor should be applied: 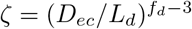 (Fig. S5).

**Fig. 3:**
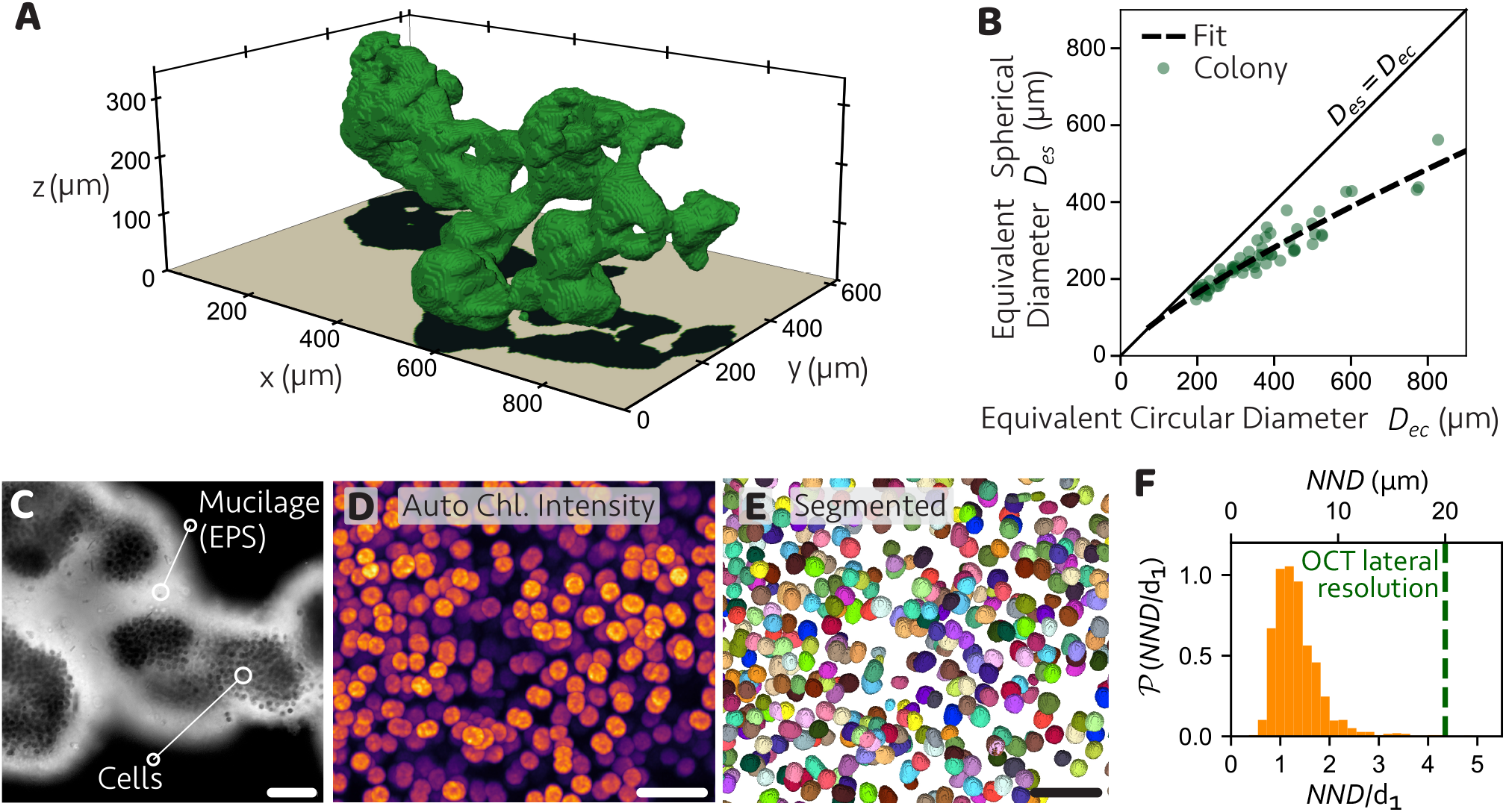
Characterization of the morphology of field samples of *Microcystis* spp. colonies. (A) Three-dimensional optical coherence tomography (OCT) scan of the colony envelope containing the cells and mucilage. (B) The projection in the imaging axis (*z*) overestimates the colony biovolume, as observed by the relation between the equivalent spherical and circular diameters measured with OCT for individual colonies of Lake’t Joppe (symbols, *N* = 66 colonies). Dashed line is the best power law fit. (C) A mucilage layer stained with Alcian Blue 8GX, visible as bright regions, encases the colony and fills the spacing between cells. Scale bar = 50 µm. (D) The three-dimensional cell arrangement within a colony was visible with confocal microscopy. Brightness intensity indicates the auto-fluorescence of chlorophyll *a*. Scale bar = 20 µm. (E) The same colony region as in panel D after the segmentation and labeling of cells using StarDist [42]. Scale bar = 20 µm. (F) Distribution of the nearest-neighbor distances (NND) within one colony normalized by the average cell diameter *d*_1_ in the bottom axis, and in absolute values in the top axis (*N* = 1404 cells). Vertical dashed line indicates the lateral resolution of the OCT scans.

We complemented the OCT measurements with mucilage staining and confocal microscopy to resolve the mucilage and cellular arrangement at the colony microscale. Staining with Alcian Blue 8GX revealed that the mucilage fills the intercellular space and forms a thin external sheath, up to a few cell diameters thick (Fig. 3C). An irregular intercellular arrangement could be observed from the confocal microscopy imaging of a few cell layers (Fig. 3D) and the subsequent segmentation of individual cells (Fig. 3E). This intercellular arrangement was quantified by a broad distribution of the normalized nearest-neighbor distance P(NND*/ d*_1_) (mean NND*/ d*_1_ = 1.35, SD = 0.55, *N* = 1404 cells, Fig. 3F). The OCT scans had a lateral resolution larger than the intercellular spacing (vertical line in Fig. 3F). Therefore, an OCT scan detected the merged volume of the cells together with the mucilage.

Besides the microstructure of the colonies, another important factor determining the buoyancy is the gas vesicle content of the cells. Here, using a capillary compression assay, we measured the average per-cell gas fraction *ϕ*_*g,c*_ in samples from Lake’t Joppe (JP: mean = 0.055, SD = 0.007, *N* = 3 replicates) and Lake Braassemermeer (BM: mean = 0.044, SD = 0.002, *N* = 3 replicates). Despite this difference in fraction, the absolute gas-volume per cell *V*_*g*_ was similar in the two lakes (JP: 2.00± 0.25 µ m^3^; BM: 2.25 ±0.12 µ m^3^), consistent with slightly larger mean cell diameters in Lake Braassemermeer (BM: mean = 4.6 µm, SD = 1.3 µm, *N* = 10285 cells) than in Lake’t Joppe (JP: mean = 4.1 µm, SD = 0.9 µm, *N* = 2168 cells). The per-cell gas fraction, together with intercellular packing and the near-water density of mucilage, determines the effective colony density (*ρ*_*a*_ in Eq.1), which directly affects the vertical velocity of the colonies.

### 3.2. Flotation velocity is influenced by colony density and size-dependent shape factors

Building on the morphological characterization, we investigated the vertical motion of individual *Microcystis* spp. colonies by particle tracking in a thermally stable density gradient (Fig. 2). A good correlation was observed between the log-transformed values of the shape factor and the equivalent circular diameter (Pearson correlation: *r*(165) = 0.87, *p <* 0.001, Fig. 4A). Small colonies (*D*_*ec*_ ≲ 100 µm) exhibited *ψ*≃ 1, whereas larger colonies reached *ψ* = 10.44 at *D*_*ec*_ = 971 µm. The dependence of *ψ* on *D*_*ec*_ follows a power law, 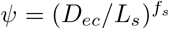, with *f*_*s*_ = 0.87 ±0.03 and *L*_*s*_ = 67 ±3 µm. This result indicates that the effect of colony morphology on the flotation velocity can be characterized by colony size, which enters Eq. (1) explicitly via the equivalent circular diameter *D*_*ec*_, but also implicitly via the shape factor *ψ*(*D*_*ec*_). Notably, the characteristic colony size *L*_*s*_ for the shape factor *ψ* is in agreement with the characteristic colony size *L*_*d*_ for the volume factor *ζ*, although the two parameters are estimated from separate datasets. This agreement is expected, as both characteristic sizes represent the same physical quantity, *i*.*e*., the colony diameter from which its shape departs from a sphere.

**Fig. 4:**
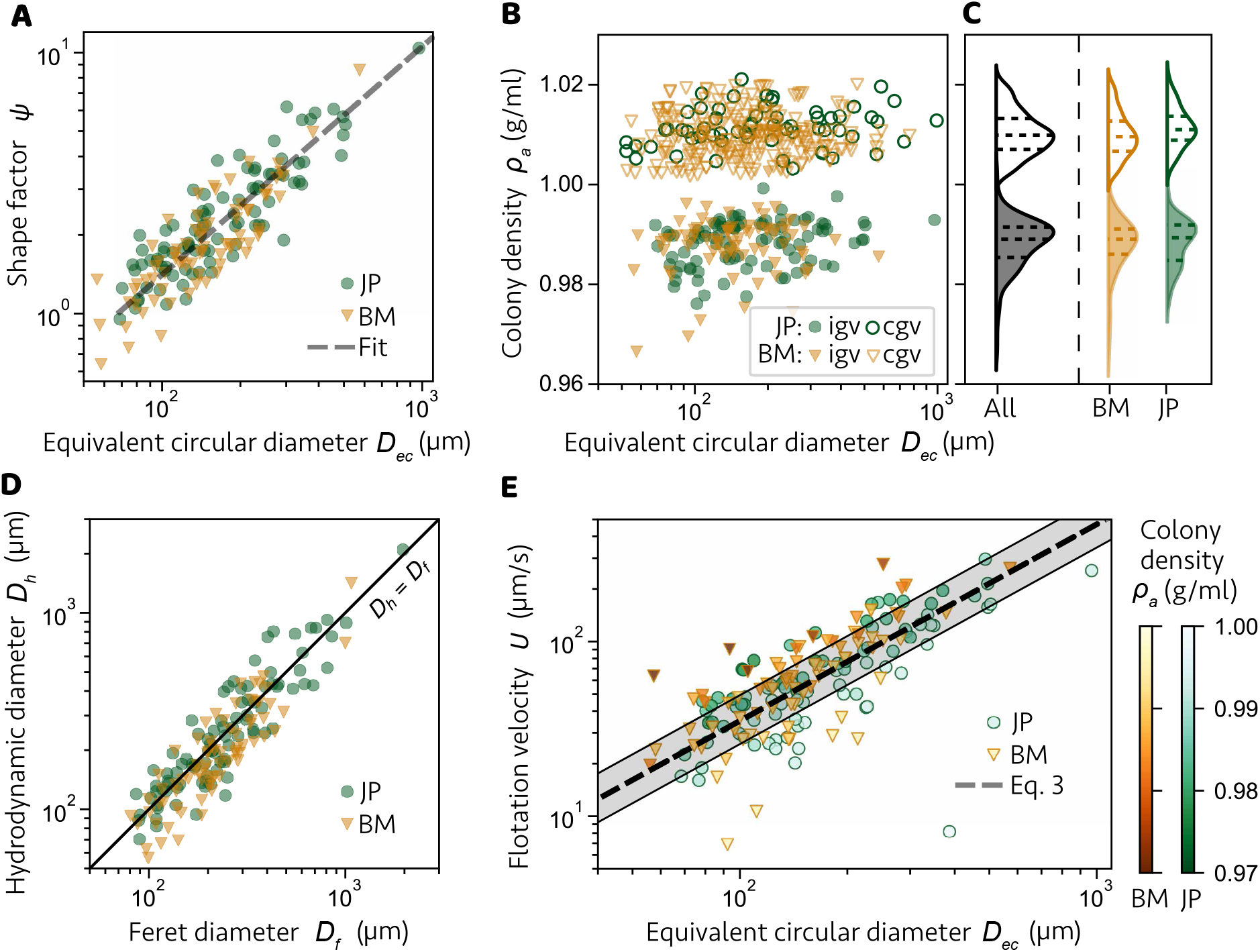
Hydrodynamics of individual *Microcystis* colonies. (A) Shape factor of individual colonies (symbols) as a function of the equivalent circular diameter. The colonies are from Lake’t Joppe (JP) and Lake Braassemermeer (BM). Dashed line is the best fit power law. (B) Density of individual colonies with intact gas vesicles (igv, filled symbols) and collapsed gas vesicles (cgv, empty symbols) as a function of the equivalent circular diameter. (C) Distribution of colony density for all samples and for each lake, separated by intact (igv, filled curve) or collapsed (cgv, empty curve) gas vesicles. Dashed lines indicate the 25^th^, 50^th^ and 75^th^ percentiles. (D) Hydrodynamic diameter of individual colonies as a function of their Feret diameter (the maximum linear dimension). (E) Flotation velocity of individual colonies in a homogeneous water column as a function of their equivalent circular diameter. Color intensity of symbols indicates the colony density. The dashed line indicates the modified Stokes’ law (Eq. 3) for the median buoyant density of the colonies, while the shaded region is bounded by the 25^th^ and 75^th^ density percentiles. The graphs show 92 colonies with intact gas vesicles from Lake’t Joppe and 75 colonies from Lake Braassemermeer. In addition, panel B contains 66 and 236 colonies with collapsed gas vesicles from Lake’t Joppe and Lake Braassemermeer, respectively.

The observed increase of the shape factor *ψ* with *D*_*ec*_ reflects systematic morphological differences between small and large colonies. To quantify these differences, we combined two corrections to the projection-based sizing. First, projection along the imaging axis overestimates buoyant force for large, branched or porous colonies; OCT quantified this bias via the volume factor *ζ* (Fig. S5). Second, hydrodynamic drag was governed by an effective hydrodynamic diameter *D*_*h*_ — the diameter of a sphere experiencing the same drag force at the same speed — which relates to the projection-based size through *D*_*h*_ = *ψ ζ D*_*ec*_ (Supporting Information Text S1). Consistent with this picture, *D*_*h*_ closely matches the Feret diameter *D*_*f*_ (maximum linear dimension) for floating colonies (Pearson correlation: *r*(165) = 0.92, *p <* 0.001) with *R*^2^ = 0.85 for the *D*_*h*_ = *D*_*f*_ curve (Fig. 4D). Thus, *D*_*f*_ serves as an appropriate size descriptor for estimating drag on *Microcystis* colonies. In turn, the large values of *ψ* observed for large colonies arise from a combined bias: projection imaging overestimates buoyant force (*D*_*ec*_ *> D*_*es*_, Fig. S5) while it underestimates drag (*D*_*ec*_ *< D*_*h*_, Fig. S6) when used as the sole size measure.

Whilst colony morphology influences velocity via its size and shape, due to the size-dependent shape factor *ψ*(*D*_*ec*_), the colony composition contributes to velocity via its density *ρ*_*a*_ (Eq. 1). In contrast to the shape factor, the density of colonies with intact gas vesicles showed only a weak but significant dependence on colony size (Pearson correlation: *r*(165) = 0.16, *p* = 0.03 for the combined datasets; Fig. 4B). Because the mucilage layer has a density close to that of water [20], colony buoyancy is dominated by the density of the cellular phase, weighted by the cell–to–colony volume fraction *ϕ*. The weak size dependence of *ρ*_*a*_ therefore suggests that the relative volumes of cells and mucilage did not vary systematically with colony size. Consistent with this, the density of colonies from the two lakes did not differ significantly (Mann-Whitney U test, *U* = 3566, *N* = 167, *p* = 0.71).

To further validate our density estimates, we collapsed the gas vesicles of the dark-incubated colonies and measured their density. The density of colonies with collapsed gas vesicles did not vary significantly with colony size (Pearson correlation: *r*(320) = 0.06, *p* = 0.31 for the combined datasets; Fig. 4B), and the variances of the intact and collapsed distributions did not differ significantly (Brown–Forsythe test, *F* (1, 467) = 1.0, *p* = 0.32; Fig. 4C), supporting the robustness of our protocol for single-colony density measurements. These constraints on *ρ*_*a*_ justify treating the density of the colonies as only weakly size-dependent over the observed range and support the decoupling of morphology (via the shape factor *ψ*) from colony density in the velocity analysis. The mean colony density with intact vesicles was 0.988 ± 0.005 g/mL (SD, *N* =167) and increased to 1.010 ±0.004 g/mL (SD, *N* = 302) after gas vesicle collapse. The difference Δ *ρ*_*a*_ between these means reflects water ingress into cells under high pressure, from which we can estimate the per-colony gas fraction, *ϕ*_*g,a*_ = 0.022 ± 0.010 (gas volume per total colony volume, i.e., cells plus mucilage). Combined with the per-cell gas fraction, *ϕ*_*g,c*_ = 0.050 (average of the two lakes), the cell-to-colony volume fraction can be estimated as *ϕ* = *ϕ*_*g,a*_*/ϕ*_*g,c*_ = 0.44 ± 0.20.

From Eq. (1), the flotation velocity in a homogeneous and quiescent water column can be calculated for each *Microcystis* colony with intact gas vesicles. The measured shape factor, density and equivalent circular diameter of each colony were the input parameters. A density of 0.9982 g/mL and a dynamic viscosity of 1.0016 mPa s were considered for water at 20 °C. The 5 % fastest floating colonies in our experiments had a velocity above 181 µm/s (15.6 m/day), while the 5 % slowest colonies were below 19 µm/s (1.6 m/day). The power law relation between the shape factor and the equivalent circular diameter, 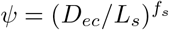, allows us to rewrite Stokes’ law for the flotation velocity of *Microcystis* spp. colonies as

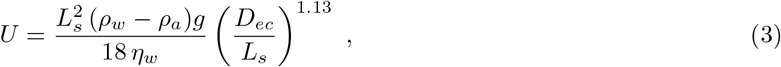

which is valid for *D*_*ec*_ ≥ *L*_*s*_, where *L*_*s*_ = 67 ± 3 µm. As expected, the flotation velocity increased with the equivalent circular diameter (Fig. 4E). However, the exponent 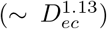 is much lower than the standard exponent for homogeneous spheres 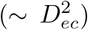, which is a result of the large shape factor measured for larger colonies (Fig. 4A). Colonies smaller than the characteristic size, *D*_*ec*_ *< L*_*s*_, were outside the measuring range of our setup. However, at *D*_*ec*_ = *L*_*s*_ the power law for the shape factor reduces to *ψ* = 1. Moreover, the shape factor of a single cell is also unitary due to its spherical shape. Therefore, we expect small colonies with *D*_*ec*_ *< L*_*s*_ to follow the classic Stokes’ law, *i*.*e*., with *ψ* = 1. Alternatively, to the equivalent circular diameter, the dependence of the flotation velocity is also given as a function of the Feret diameter of colony (Supporting Information Text S1, Fig. S6), as the latter is a convenient size descriptor for microscopy.

Our findings deviate from the common assumption that vertical velocity varies with the square of colony size [13, 14, 21]. Instead, our results agree with previous studies that found an increase in the shape factor with colony size [22, 23]. Notably, the exponent obtained here 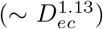 is lower than that found by Nakamura *et al*. [22] 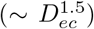. However, Nakamura *et al*. [22] defined the characteristic size *L*_*s*_ as the distance between neighboring cells (7.2 *µ*m in their work), while here we allow the characteristic size to be a fitting parameter. Setting the characteristic length to be the average nearest-neighbor distance for our data (6.2 *µ*m, Fig. 3F) and recalculating the power law fit for our measured flotation velocities, we obtained a higher exponent 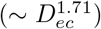, though with a lower coefficient of determination (*R*^2^ = 0.39). Differences between our results and those of Nakamura *et al*. [22] might be due to differences in morphospecies composition, which were dominated by *M. ichthyoblabe* in our samples but not identified in Nakamura *et al*. [22]. Alternatively, Li *et al*. [23] fitted the dependence of the shape factor with colony size using a second-order polynomial. The different functional form of the fit does not allow for a direct comparison with our model parameters. However, the colony sizes and measured flotation velocities in Li *et al*. [23] were on average larger than the values from our experiments. Overall, the larger sample size and the use of two different lakes provide a higher confidence level for our results compared to previous measurements.

In line with many previous studies, we demonstrate that the colony density has a large impact on the velocity and can explain much of the variation observed in our experiments (Eq. 3, Fig. 4E). That is, the velocity ratio between the 10^th^ and 90^th^ percentiles of colony density 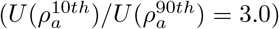 is almost of the same order of magnitude as the velocity ratio between the 90^th^ and 10^th^ percentiles of colony size 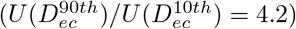. Furthermore, the dependence of the shape factor on colony size observed in our measurements (Fig. 4A) disfavors the use of a fixed shape factor across sizes to estimate the buoyant density, as applied in previous studies [23, 26].

The large effect of colony shape and density on its vertical velocity also draws attention to morphological variation among morphospecies of *Microcystis*, which differ in outline, cell packing, and extracellular matrix composition of the colonies. Furthermore, differences in growth conditions are also known to affect colony morphology and EPS production [48, 49]. Variation in the mucilage volume fraction can have major consequences for the vertical velocity of colonies. Mucilage has a density close to water and therefore contributes little to buoyancy [20]. Instead, increased mucilage enlarges the effective size and projected area of a colony, raising viscous resistance, which slows its vertical velocity.

### 3.3. Ecological implications

Size-dependent variation of the shape factor affects the vertical migration patterns of *Microcystis* colonies in the water column. We exemplified this effect by applying the classic Stokes’ law (Eq. 1) and our modified Stokes’ law (Eq. 3) to a standard vertical migration model of *Microcystis* colonies [6, 7, 10]. The model predicts the vertical motion of a colony in a quiescent water column, assuming that the density of the colony is regulated by photosynthetic production and respiratory consumption of carbohydrates. The carbohydrate content of the cells increases when photosynthesis exceeds respiration, *i*.*e*., when the colonies are exposed to favorable light conditions in the upper part of the water column at daytime. Accumulation of carbohydrates increases the density of the colonies and they start sinking. Conversely, the carbohydrate content decreases by respiration when colonies are deprived of light deeper down in the water column and at night. Loss of carbohydrates reduces the density of the colonies, and they float upwards again. Hence, the colonies cycle up and down through the water column.

Colony size affects the vertical velocity of colonies and hence their migration patterns, as seen in a bifurcation plot displaying the local minimum and maximum depths of the colony trajectories as a function of colony size (Fig. 5A). Under the classic Stokes’ law (*ψ* = 1 in Eq. 1), small colonies develop multi-periodic or aperiodic migration trajectories indicative of chaotic dynamics, as characterized by a large spread in the local minima and maxima in the bifurcation diagram for colony diameters of 20 *< D*_*ec*_ *<* 100 µm and 170 *< D*_*ec*_ *<* 260 µm (upper panel in Fig. 5A). Similar chaotic trajectories have been described in earlier model studies of the vertical migration of cyanobacterial colonies [10, 11, 50]. Colonies with diameters of 110 *< D*_*ec*_ *<* 150 µm have periodic migration, though with a period of 2 days (Fig. S7). Larger colonies of *D*_*ec*_ *>* 260 µm develop periodic migration patterns with one or more migration cycles per day. Our modified Stokes’ law (Eq. 3) predicts slower vertical velocities due to the increase of the shape factor for colonies larger than *D*_*ec*_ *> L*_*s*_, keeping all other parameters unchanged. Notably, this implies a much wider range of sizes of colonies displaying multi-periodic or aperiodic migration, where periodic migration with one or more cycles per day is only achieved for colonies of *D*_*ec*_ *>* 710 µm (lower panel in Fig. 5A). These different migration patterns can also be observed from time plots (Fig. 5B) and phase portraits (Fig. 5C) of three representative colony sizes. For a small colony of *D*_*ec*_ = 50 µm (below *L*_*s*_), the modified Stokes’ law reduces to the classic version, and migration trajectories show chaotic patterns that do not synchronize with the day-night cycle. Meanwhile, a moderately sized colony of *D*_*ec*_ = 350 µm will display a periodic daily vertical migration under the classic Stokes’ law, but chaotic vertical migration under the modified Stokes’ law. Lastly, a very large colony of *D*_*ec*_ = 720 µm will synchronize its migration with the day-night cycle in both models, though the classic Stokes’ law leads to two migration cycles per day (Fig. 5B) and a small amplitude of the density variation (Fig. 5C), in contrast with a single daily migration cycle and larger density amplitude for the modified Stokes’ law.

**Fig. 5:**
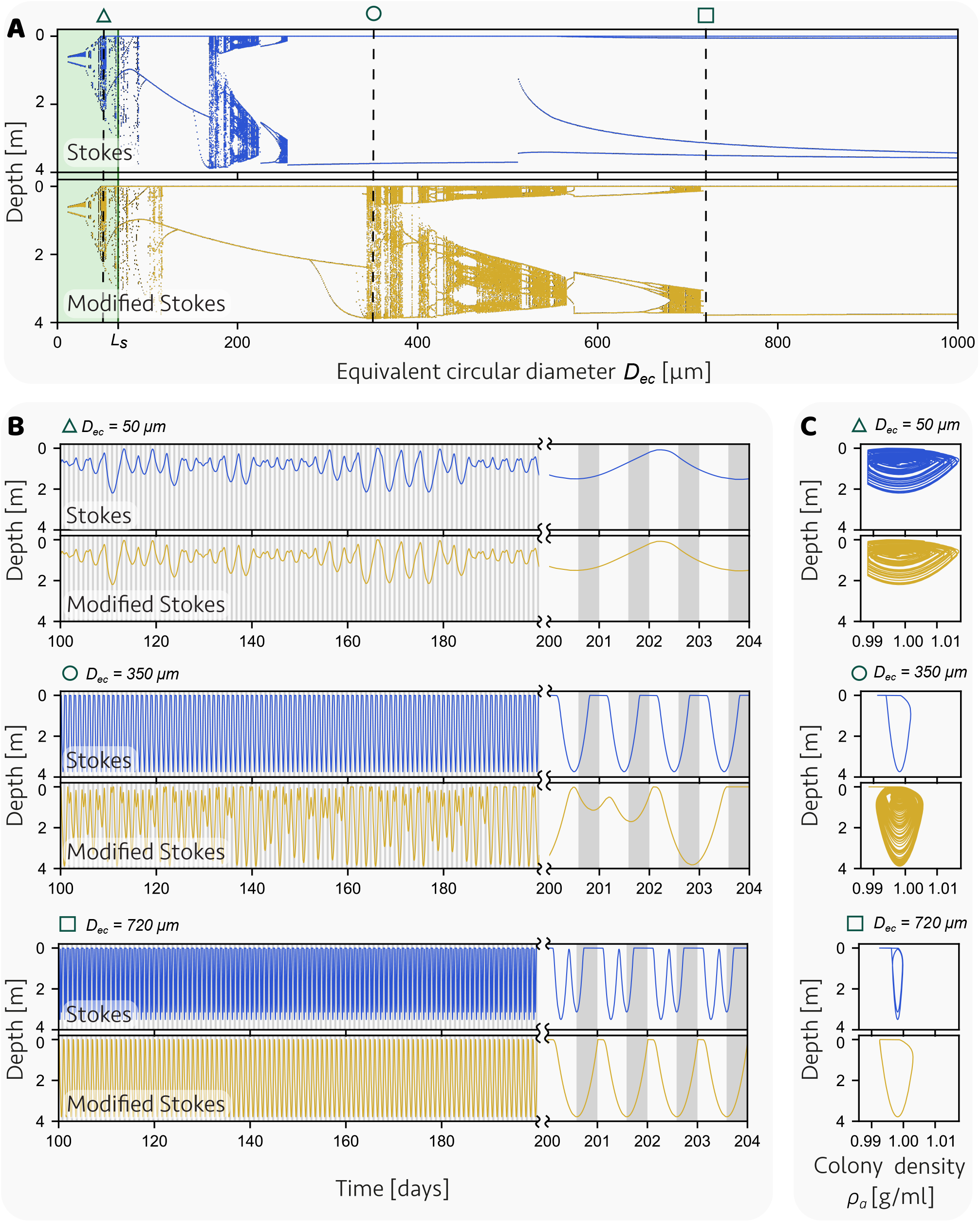
Simulations of the vertical migration of *Microcystis* colonies as modeled in Feng *et al*. [10] using the classic Stokes’ law (Eq. 1 with *ψ* = 1) or the modified Stokes’ law (Eq. 3). (A) Bifurcation plots depicting the local minimum and maximum depths of a colony between days 50 and 250, as a function of the equivalent circular diameter. Shaded green region indicates size range for which the modified Stokes’ law reduces to the classic version (*ψ* (*D*_*ec*_ ≤*L*_*s*_) = 1). (B) Depth of a colony as a function of time for three representative colony sizes (*D*_*ec*_ = 50, 350 and 720 µm). Note the shift in time scale at day 200, to provide a more detailed view of the daily patterns. Vertical shaded regions indicate periods of darkness. (C) Corresponding trajectories of the colony plotted in the phase plane of colony density and depth for three representative colony sizes (*D*_*ec*_ = 50, 350 and 720 µm). Values for model parameters are listed in the Supplementary Table S2.

These new insights can be readily used in models that aim to predict cyanobacterial blooms. For example, since our data are based on individual colonies, the new scaling of vertical velocity with colony size is well suited to be implemented in Individual-Based Models (IBMs) to predict colony-level variability and population dynamics [12]. In addition, accurate prediction of the flotation velocity of *Microcystis* colonies is also relevant for the design of artificial deep-mixing systems aiming to control bloom formation in lakes and reservoirs [15]. For an effective design, the enhanced turbulence induced by artificial mixing must be high enough to entrain the fast-floating colonies, so that they cannot float upwards to form a dense bloom at the water surface [5]. However, overestimates of the vertical velocity of colonies could lead to excessive mixing and hence excessive energy costs of artificial mixing, *e*.*g*., when bubble plumes are used as a mixing device. Therefore, the results and data provided here may help with the accuracy of prediction models used for the optimal design of artificial mixing in lakes [15].

Future studies could compare the shape factors of distinct morphospecies of *Microcystis* or other species of colonial cyanobacteria. Furthermore, high particle concentrations can lead to hydrodynamic interactions between these particles [51], and hence further study of the flotation of colonies in dense cyanobacterial suspensions is desirable. Beyond cyanobacteria, our methodology can be applied to other irregularly shaped aggregates in aquatic ecosystems (e.g., marine snow, algae–microplastic hetero-aggregates), enabling independent measurements of size, shape, and density to improve vertical-transport predictions. In addition to these ecological applications, our techniques to measure the flotation velocity, density and shape factor of a large number of individual cell aggregates may also provide interesting tools for, e.g., photo-bioreactor design and wastewater management.

## 4. CONCLUSIONS

This study advances current knowledge on the flotation velocity of the cyanobacterium *Microcystis* by providing accurate measurements of the shape factor and buoyant density at single-colony resolution. We demonstrate that the shape of colonies varies systematically with their size. As a consequence, the vertical velocity of colonies does not scale with the square of colony size but with an exponent considerably lower than predicted by classic Stokes’ law. Moreover, we show that the hydrodynamic drag of colonies can be estimated from their maximum linear dimension, a quantity easily accessible via microscopy. Implementation of the new size-dependent shape factor in a vertical migration model predicts slower migration of large colonies, thus extending the range of colony sizes that display complex migration patterns and restricting large colonies to one migration cycle per day. Our results show that consideration of colony size and shape improves understanding of the vertical migration of cyanobacterial colonies, which plays a key role in cyanobacterial bloom formation.

## ACKNOWLEDGMENTS

We thank the van Leeuwenhoek Centre for Advanced Microscopy (LCAM-FNWI) for the support with confocal microscopy. We are grateful to Pieter Slot, Merijn Schuurmans, and Axel Gunderson for support with fieldwork and sample incubation, Erik Hop for support with the experimental design, Micol Cresto-Dina for assistance with the setup calibration, Nico Schramma for support with image processing, and Isaac Deen Garcia for assistance with the OCT scans. We especially thank Johan Oosterbaan and Richard Steel for their contributions during the project’s conception. We acknowledge funding from the Hoogheemraadschap van Rijnland. MJ acknowledges support from the ERC Starting Grant “2023-StG-101117025, FluMAB.

## DATA AND CODE AVAILABILITY

Primary data of the colony tracking experiments are available at https://doi.org/10.5281/zenodo.18959105 and primary data for the colonial morphology characterization are available at https://doi.org/10.5281/zenodo.18959341. The scripts for image analysis and the vertical migration model are available at https://doi.org/10.21942/uva.31650862

## SUPPORTING INFORMATION FOR

### Supporting Information Text

#### S1. Buoyancy-driven motion of non-spherical particles

Consider a colony with total volume *V*_*c*_ (cells and mucilage) and colony density *ρ*_*a*_ floating in a homogeneous and quiescent liquid column of density *ρ*_*l*_. The net buoyant force on the colony is

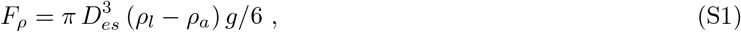

where *g* is the gravity acceleration, and *D*_*es*_ = (6 *V*_*c*_*/ π*)^1/3^ is the equivalent spherical diameter. A positive force value indicates an upwards force (*i*.*e*., opposite to gravity). Let *U* be the rising velocity of the colony. The viscous drag force *F*_*h*_ on the colony is

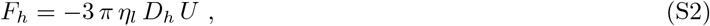

where *η*_*l*_ is the liquid dynamic viscosity and *D*_*h*_ is the hydrodynamic diameter of the colony, *i*.*e*., the diameter of a sphere with equal drag force under the same rising velocity. Here we assume that the Reynolds number, *Re* = *ρ*_*l*_ *U D*_*h*_*/ η*_*l*_, is sufficiently small such that inertial effects may be neglected. The balance between the buoyant force (Eq. S1) and viscous drag (Eq. S2) results in the modified Stokes’ law (1), where the shape factor is defined as *ψ* = *D*_*h*_*/ ζ D*_*ec*_ and the volume factor is defined as *ζ* = (*D*_*es*_*/ D*_*ec*_)^3^. The power-law dependence of the volume factor, 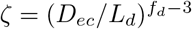, and shape factor, 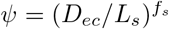, on the equivalent circular diameter can be extended to the hydrodynamic diameter, such that, 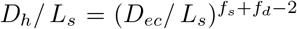, taking *L*_*d*_ = *L*_*s*_ (Fig. S6A). Consequently, the modified Stokes’ law can also be written in terms of the hydrodynamic diameter,

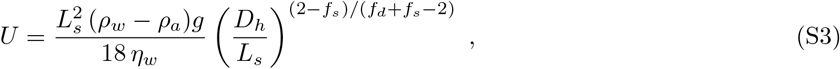

where the hydrodynamic diameter can be approximated by the Feret diameter *D*_*h*_ = *D*_*f*_ (Fig. S6B).

The force balance on the colony is modified if the liquid column has a linear density gradient *ρ*_*l*_(*z*) = *ρ*_*o*_ − *β z*, where *z* is the height from the bottom of the chamber, *ρ*_*o*_ is the density at the bottom and *β* is a constant. In this case, the rising colony will drag denser fluid into lighter layers and therefore distort the isopycnic surfaces. The added weight of the denser fluids modifies the vorticity field around the colony, thus enhancing the drag force [1]. For a spherical particle in a non-diffusive liquid, the ratio *K*_*e*_ = *F*_*e*_*/F*_*h*_ between the entrainment drag *F*_*e*_ and homogeneous drag *F*_*h*_ was obtained by Yick *et al*. [2], given by

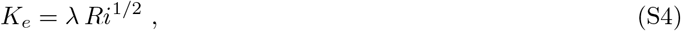

where 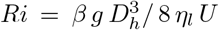 is the viscous Richardson number and *λ* = 1.95 is a numerical pre-factor. The non-diffusive limit is characterized by a large Péclet number, *i*.*e*., *Pe* = *U D*_*h*_ /2 *κ* ≫1, where *κ* is the Percoll diffusivity. Here, we extend the correction from Yick *et al*. [2] to non-spherical colonies by substituting the sphere diameter with the hydrodynamic diameter of the colony, which is the length scale that governs the flow disturbance around the colony. We verify this assumption by comparing the flotation velocities obtained in the linear density gradient with the values obtained in a discrete double layer liquid column (Fig. S4). The entrainment and homogeneous drag forces must balance the buoyant force, resulting in an implicit expression for the rising velocity of the colony as a function of the height, *U* (*z*), given by

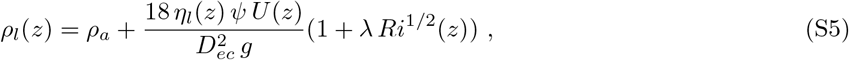

where *Ri*(*z*) = *β g* (*ψ ζ D*_*ec*_)^3^/ 8 *η*_*l*_(*z*) *U* (*z*).

#### S2. Comparison of the flotation velocity measured in a discrete double layer and in a linear density gradient

The implicit expression for the flotation velocity, Eq. 2 requires as a minimum of two pairs of input values, namely the colony velocity *U* and liquid density *ρ*_*l*_, to extract two output values, namely the shape factor *ψ* and the colony density *ρ*_*a*_. A linear density gradient provides a large range of pairs of input values due to continuously varying density. However, a column with two discrete layers can potentially be used to extract the output parameters. Therefore, we compared the measured flotation velocity of colonies using the linear density gradient (Fig. S4A) and an alternative column with a discrete double layer of Percoll (Fig. S4B). While the first liquid column produces a thermally stable column (Fig. S4A.i) due to the linear density profile (Fig. S4A.ii), the discrete double layer has thermal convection currents (Fig. S4B.i) in each of the homogeneous layers (Fig. S4B.ii). While rising colonies in the linear density gradient have negligible horizontal velocities (Fig. S4A.iii and iv), the colonies in the double layer have both horizontal and vertical velocity bias (Fig. S4B.iii and iv). Furthermore the velocity greatly decreases near the sharp interface of the double layer due to the intense fluid entrainment [3]. Consequently, many colony trajectories in the double layer are lost due to horizontal drift out of the field of view, or overlap in the sharp interface. Moreover, the vertical velocity bias (∼22 µm/s) in the bulk of each layer must be corrected using the reference trajectories of calibrated microspheres. After correction of the velocity bias, the two methodologies provide a good agreement in the measured flotation velocities (Fig. S4C). One-way Analysis of Covariance test was performed between the two techniques, with the log-transformed value of flotation velocity as dependent variable and the log-transformed value of equivalent circular diameter as covariate, without statistically significant differences (ANCOVA, *F* (1, 142) = 0.29, *p* = 0.6). However, the linear density gradient is more suitable for velocity measurements as many colony trajectories in the discrete double layer extended beyond the field of view (Fig. S4B.iii) and thus could not be fully tracked.

#### S3. Change in cell density after the collapse of gas vesicles

The process of collapse of gas vesicles inside the cells of buoyant cyanobacteria has been reviewed by Walsby [4]. Let *V*_1_ be the volume of a single cell and *V*_*g*_ be inner volume of the gas vesicles inside the cell. The gas vesicles will fracture When high pressure is applied (1 MPa is sufficient to collapse all gas vesicles in *Microcystis* cells [4]). Immediately, the gas will dissolve and the cell will contract by a volume *V*_*g*_. Consequently, the turgor pressure inside the cell will drop and water will enter the cell to balance the difference in solute potential. The volume of water that enters the cell is Δ*V*_*w*_, thus the final volume of the cell after collapse of gas vesicles will be 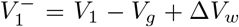. If *ρ*_*c*_ is the density of the cell with intact gas vesicles, the density of the cell after collapse of gas vesicles will be

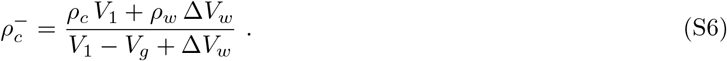

The volume of water entering has been estimated as Δ*V*_*w*_ = 0.73 *V*_*g*_ [5]. For simplicity, we approximate Δ*V*_*w*_ ≈*V*_*g*_, such that the volume of the cell remains constant after gas vesicles collapse 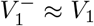, also applied and verified via electron microscopy by Nakamura *et al*. [6]. Therefore, the change in cell density before and after gas vesicle collapse is

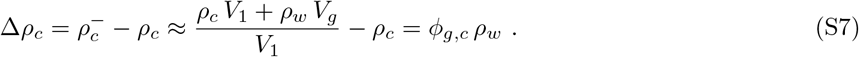

The error associated with the constant cell volume approximation in the collapse cell density is 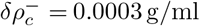, which is below our instrumental uncertainty in density measurement (*δ*_*ins*_*ρ* = 0.0006 g/ml) and thus negligible.

## Supplementary Tables

**Table S1:** Morphological descriptors for *Microcystis* colonies.

| Name | Symbol | Definition |
| --- | --- | --- |
| **Descriptors based on hydrodynamics** |  |  |
| Hydrodynamic diameter | $D_h$ | Diameter of a sphere with equal Stokes drag force |
| Shape factor | $\psi = D_h / \zeta D_{ec}$ | Ratio between the hydrodynamic diameter and a characteristic length, chosen here to be the equivalent circular diameter |
| **Descriptors based on composition/structure in 3D region** |  |  |
| Colony volume | $V_c = V_1 N_c + V_m$ | Volume occupied by cells and mucilage |
| Cell-to-colony volume fraction | $\phi = V_1 N_c / V_c$ | Ratio between total volume of cells per total colony volume (cell+mucilage) |
| Colony density | $\rho_a = \rho_c \phi + \rho_m (1 - \phi)$ | Density weighted by the cell and mucilage proportions |
| Equivalent spherical diameter | $D_{es} = (6V_c / \pi)^{1/3}$ | Diameter of a sphere with equal volume |
| Average nearest-neighbor distance | $\overline{\text{NND}}$ | Average value of the distributions of nearest-neighbor of each cell |
| **Descriptors based on 2D projection** |  |  |
| Projected area | $A_c$ | Projection of the 3D region along a given axis |
| Feret diameter | $D_f$ | Maximum distance between any two points of the colony in a 2D projection |
| Equivalent circular diameter | $D_{ec} = (4 A_c / \pi)^{1/2}$ | Diameter of a circle with same projected area |
| Major/minor axis | $a, b$ | Major/minor axis of an ellipse with the same second order moments of the projected area |
| Aspect ratio | $AR = a/b$ | Ratio between the major and minor axis |

**Table S2:** Model parameters.

| Parameter | Value |
| --- | --- |
| Cell-to-colony volume fraction | $\phi = 0.44$ |
| Density of cells without gas vesicles and carbohydrate storage | $\rho_{min} = 1.0201 \text{ g/ml}$ |
| Per-cell gas fraction (average of the two lakes) | $\phi_{g,c} = 0.050$ |
| Shape factor | (i) Classic Stokes: $\psi = 1$;<br>(ii) Modified Stokes: $\psi(D_{ec} < L_s) = 1$ or<br>$\psi(D_{ec} \geq L_s) = (D_{ec}/L_s)^{f_s}$, where<br>$L_s = 67 \mu\text{m}$ and $f_s = 0.87$ |
| Temperature | $T = 20^\circ\text{C}$ |
| Water density | $\rho_w = 0.9982 \text{ g/ml}$ |
| Water viscosity | $\eta_w = 1.0016 \text{ mPa s}$ |
| Light attenuation coefficient [7] | $K_t = 4.4 \text{ m}^{-1}$ |
| Day-night cycle [8] | L:D = 14:10 h |
| Maximum incident light intensity at water surface [8] | $I = 1000 \mu\text{mol photons m}^{-2} \text{ s}^{-1}$ |
| Minimum cellular carbohydrate content [8] | $Q_{min} = 0.2 \mu\text{mol glucose mm}^{-3}$ |
| Carbohydrate specific density [8] | $c = 29.0 \text{ kg mm}^{-3} (\mu\text{mol glucose mm}^{-3})^{-1}$ |
| Normalization constant [8] | $A = 1.275 \times 10^7 \mu\text{mol glucose mm}^{-3} \text{ d}^{-1}$ |
| Activation energy [8] | $E_A = 0.4105 \text{ eV}$ |
| Scaling parameter of photosynthetic carbohydrate production rate in the light [8] | $Y_T = 0.0687 \mu\text{mol glucose mm}^{-3} \text{ d}^{-1} (\mu\text{mol photons m}^{-2} \text{ s}^{-1})^{-1}$ |
| Optimum light intensity [8] | $I_{opt,T} = 269.2 \mu\text{mol photons m}^{-2} \text{ s}^{-1}$ |
| Carbohydrate respiration rate in the dark [8] | $R_T = 1.1232 \mu\text{mol glucose mm}^{-3} \text{ d}^{-1}$ |
| Mucilage density [9] | $\rho_m = 0.9989 \text{ g/ml}$ |
| Minimum colony density [10] | $\rho_{a,min} = 0.920 \text{ g/ml}$ |
| Maximum colony density [10] | $\rho_{a,max} = 1.065 \text{ g/ml}$ |

## Supplementary Figures

**Fig. S1:**
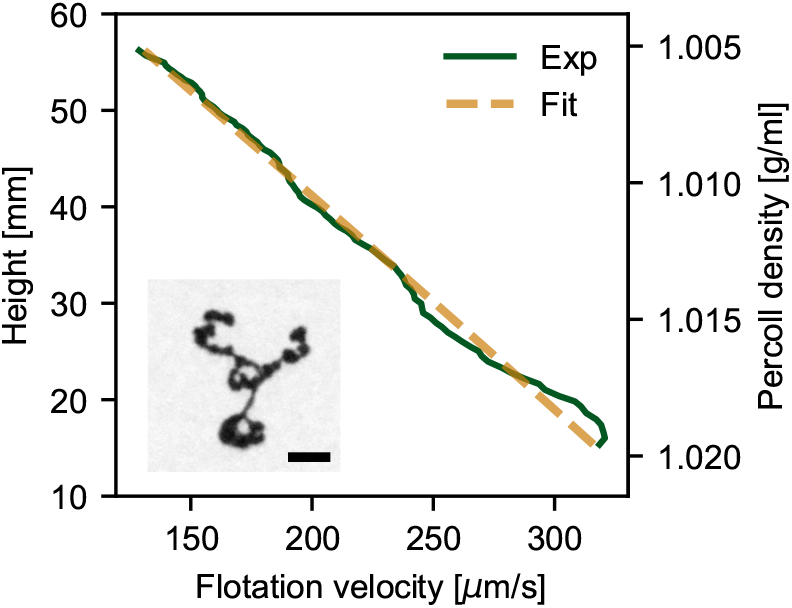
Example of velocity trajectory (solid line) and best fit (dashed line) from Eq. (2) for a single colony of *Microcystis* rising in the flotation chamber. Inset shows the image of the colony. Scale bar = 200 µm. *D*_*ec*_ = 344 µm, *ψ* = 3.1, *ρ*_*a*_ = 0.992 g/mL, *Re* = 0.045, *Ri* = 0.16, *Pe* = 3.2 · 10^3^.

**Fig. S2:**
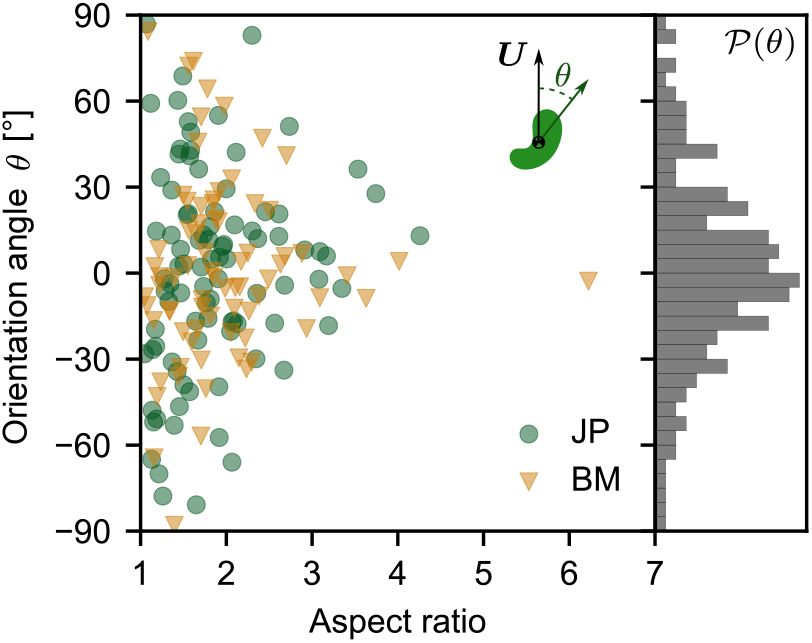
Left panel: Orientation angle between the major axis and the velocity of the colonies of *Microcystis* spp. as a function of its aspect ratio (major/minor axis). Right panel: normalized distribution of orientation angles.

**Fig. S3:**
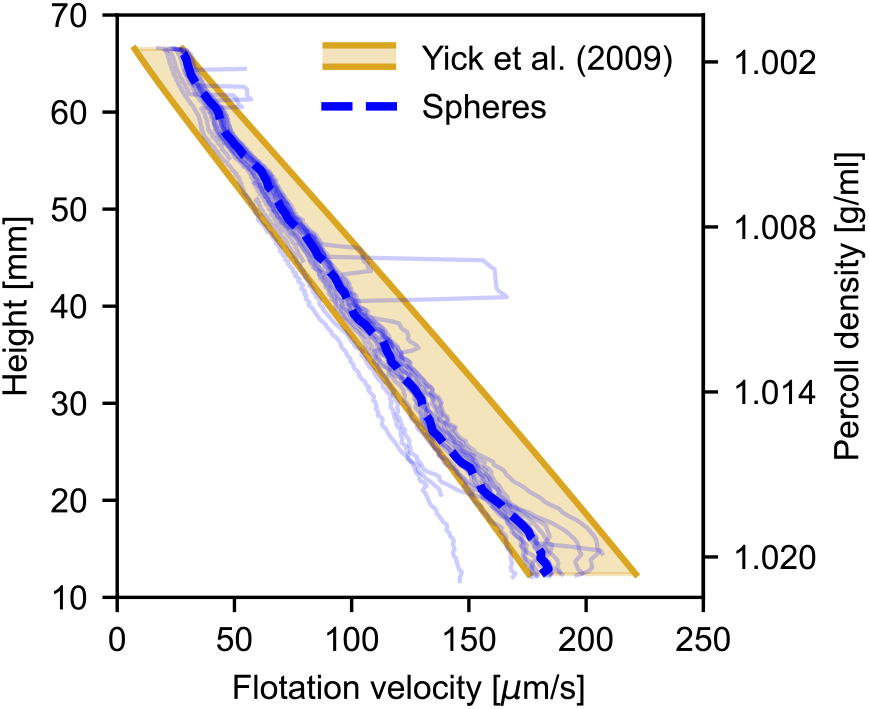
Validation of flotation chamber using calibrated spheres. Chamber is filled with a linear density gradient of a Percoll suspension. Spheres have a diameter of 148 ±5 µm (SD, *N* = 56 spheres), density between 0.9983 g/mL to 1.0003 g/mL and eccentricity *<* 0.3. Dashed line indicates the median velocity of all velocity trajectories (*N* = 20 spheres) and shaded region indicates the theoretical prediction from Eq. (2) within the uncertainty range of the diameter and density of the spheres [2].

**Fig. S4:**
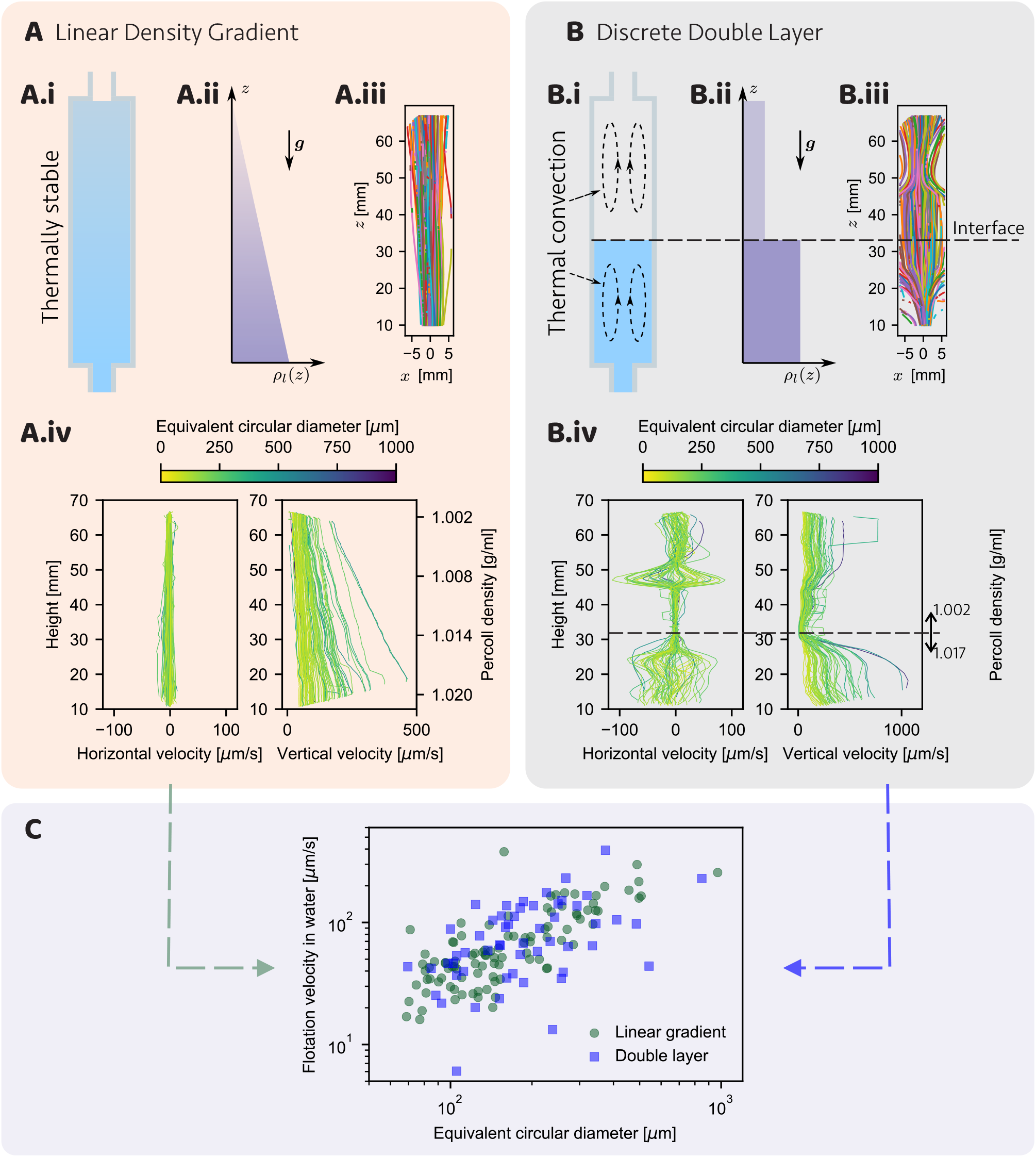
Velocity measurement of *Microcystis* spp. colonies in two liquid column structures: (A) Linear density gradient and (B) Discrete double layer. Both columns are prepared with Percoll suspension at different concentrations. (i) Diagram of liquid column. (ii) Diagram of liquid density as a function of height *z*. (iii) Colony trajectories. (iv) Horizontal (*x*) and vertical (*z*) velocities of each colony trajectory. (C) Flotation velocity of each colony in a homogeneous water column as a function of the equivalent circular diameter, measured with both liquid column structures. Sample collected at Lake’t Joppe. Number of colonies: *N* = 91 (linear density gradient) and 55 (discrete double layer).

**Fig. S5:**
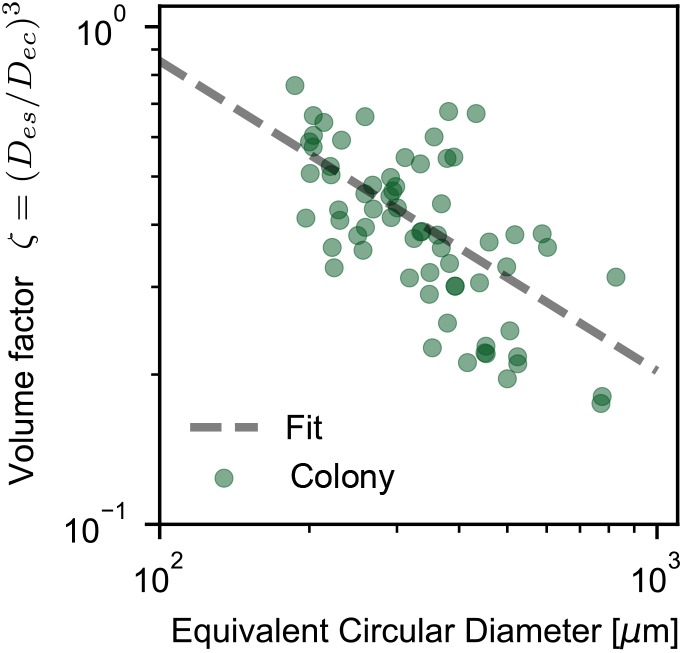
Volume factor *ζ* = (*D*_*es*_*/D*_*ec*_)^3^ as a function of the equivalent circular diameter measured with OCT scans of *Microcystis* spp. colonies collected from Lake’t Joppe (*N* = 66 colonies). Dashed line indicates the power law 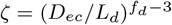, where *L*_*d*_ = 77 ± 20 µm and *f*_*d*_ = 2.36 ± 0.10.

**Fig. S6:**
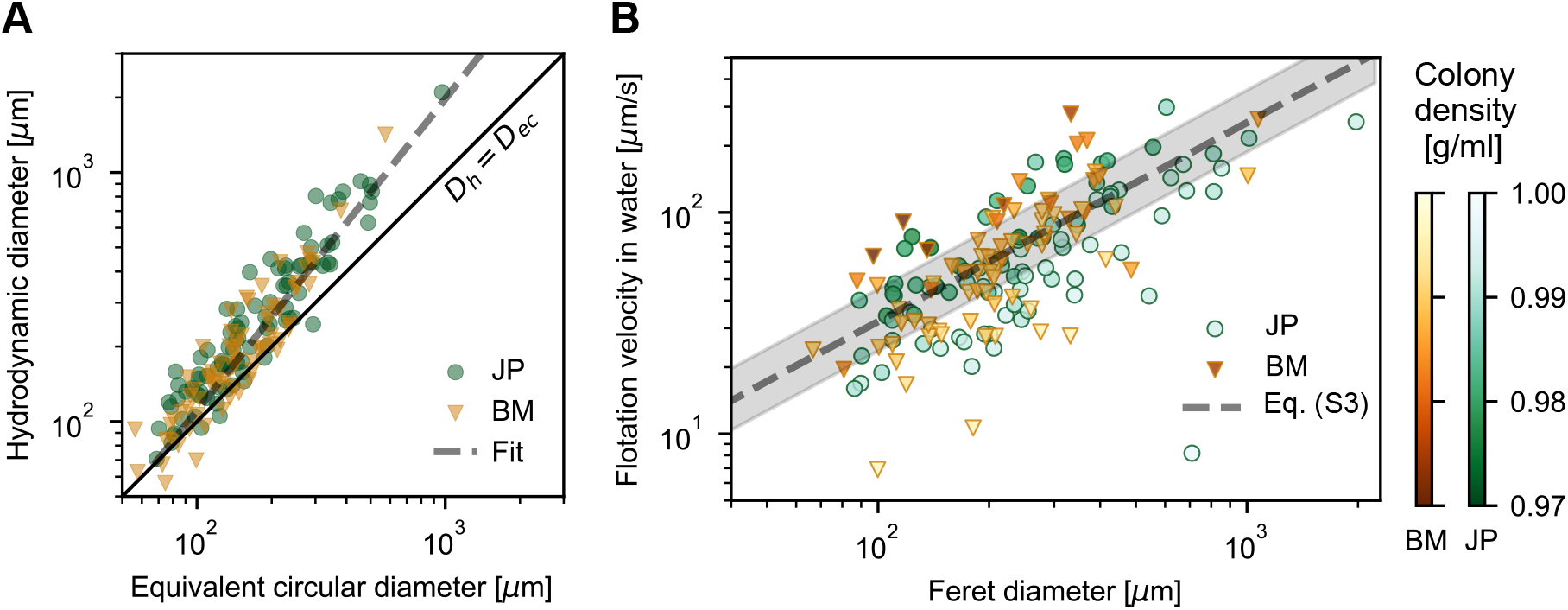
(A) Estimate of the hydrodynamic diameter (*D*_*h*_ = *ψ ζ D*_*ec*_) for each colony as a function of the equivalent circular diameter, measured from the microscopic camera in the flotation chamber. Solid line is *D*_*h*_ = *D*_*ec*_ curve. Dashed line indicates the power law 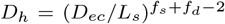, where *L*_*s*_ = 67 ±3 µm, *f*_*s*_ = 0.87 ±0.03 and *f*_*d*_ = 2.36 ±0.10. (B) Flotation velocity of individual colonies in a homogeneous water column as a function of their Feret diameter. Color intensity of symbols indicates the colony density. The dashed line indicates the modified Stokes’ law written in terms of the Feret diameter (Eq. S3, power-law dependence of 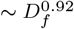,) for the median buoyant density of the colonies, while the shaded region is bounded by the 25^th^ and 75^th^ density percentiles. All panels display *N* = 167 colonies.

**Fig. S7:**
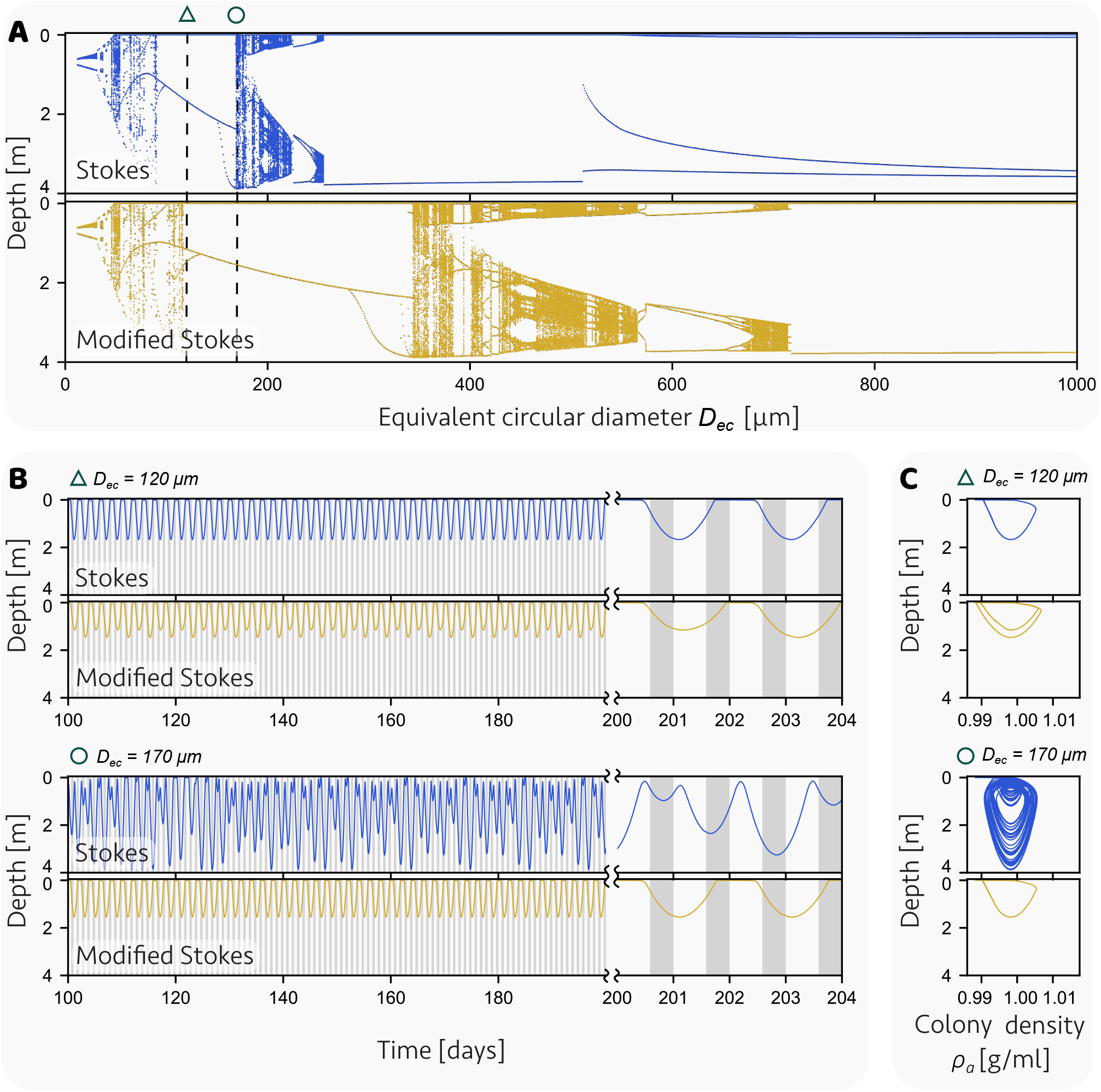
Additional plots of the simulations of the vertical migration of *Microcystis* colonies as modeled in Feng *et al*. [8] using the classic Stokes’ law (Eq. 1 with *ψ* = 1) or the modified Stokes’ law (Eq. 3). (A) Bifurcation plots depicting the local minimum and maximum depths of a colony between days 50 and 250, as a function of the equivalent circular diameter. (B) Depth of a colony as a function of time for two representative colony sizes (*D*_*ec*_ = 120, 170 µm). Note the shift in time scale at day 200, to provide a more detailed view of the daily patterns. Vertical shaded regions indicate periods of dark. (C) Corresponding trajectories of the colony plotted in the phase plane of colony density and depth for two representative colony sizes (*D*_*ec*_ = 120, 170 µm). Values for model parameters are listed in the Supplementary Table S2.

